# SARS-CoV-2 Delta variant pathogenesis and host response in Syrian hamsters

**DOI:** 10.1101/2021.07.24.453631

**Authors:** Mohandas Sreelekshmy, Yadav Pragya Dhruv, Shete Anita, Nyayanit Dimpal, Sapkal Gajanan, Lole Kavita, Gupta Nivedita

## Abstract

B.1.617 lineage is becoming a dominant SARS-CoV-2 lineage worldwide and was the dominant lineage reported in second COVID-19 wave in India, which necessitated studying the properties of the variant. We evaluated the pathogenicity and virus shedding of B.1.617.2 (Delta) and B.1.617.3 lineage of SARS-CoV-2 and compared with that of B.1, an early virus isolate with D614G mutation in Syrian hamster model. Viral load, antibody response and lung disease were studied. No significant difference in the virus shedding pattern was observed among these variants studied. A significantly high SARS-CoV-2 sub genomic RNA could be detected in the respiratory tract of hamsters infected with Delta variant for 14 days. Delta variant induced lung disease of moderate severity in 40% of infected animals. The neutralizing capability of the B.1, Delta and B.1.617.3 variant infected animals were found significantly lower with the B.1.351 (Beta variant). The findings of the study support the attributed disease severity and the increased transmission potential of the Delta variant.

## Introduction

SARS-CoV-2 B.1.617 lineage variants were first reported from India in October 2020. Among the three sub lineages of B.1.617 reported, B.1.617.1 is designated as Variant of Interest and B.1.617.2 as Variant of Concern (VOC) by World Health Organization [1]. As on 18^th^ July 2021, Delta variant has been reported in 124 countries [1]. Among the 3 sub lineages, B.1.617.3 is the variant of which only few sequences have been detected. During the initial phase of the second wave of COVID-19 in India, B.1.1.7 and B.1.617.1 were detected the most. But the trend suddenly changed and the B.1.617.2 (Delta) sub lineage increased with the surge in COVID-19 cases later. The rise in cases in India was speculated to a potential high transmission potential of this variant which replaced the other variants in the circulation like B.1.617.1 and B.1.1.7 [2]. The characteristic mutations reported in the spike gene of the B.1.617 lineage is D111D, L452R, D614G, P618R and ±E484Q [3]. These mutations suggest increased ACE2 binding, transmissibility and escape of neutralization [3–6]. The high growth rate observed in COVID-19 cases in India could be due to the combination of properties of transmissibility and immune escape.

United Kingdom has reported the largest number of cases due to B.1.617 sub lineages outside India. The risk assessment data published by the Public Health England points to higher transmissibility compared to the wild type virus, immune escape potential and hospitalization risk of the Delta variant [7]. The potential impacts of the variant on vaccine and therapeutic effectiveness is uncertain as limited data is available at present. Few recent studies have reported the neutralization efficiency of vaccinated individuals and monoclonal antibody-based therapies against Delta variant [8–12]. Although vaccination could protect against these variants, inequitable distribution of vaccines around the world can contribute to the spread. Hence more studies on the epidemiological characteristics of the emerging variants are the need of the hour, which could alert us on possible implications and for better preparedness.

In the present study we assessed the pathogenicity and virus shedding differences of the Delta and B.1.617.3 variants in Syrian hamster model, to understand whether virus shedding or tissue tropism has any role in its increased transmissibility and severity. We also investigated whether there are any differences in neutralization potential among the variants. The study parameters were compared with B.1 (an early SARS-CoV-2 isolate) variant to understand the differences of the B.1.617 with that of earlier circulating variant.

## Materials and Methods

### Virus and cells

SARS-CoV-2 variants Delta (GISAID identifier: EPI_ISL_2400521), B.1.617.3 (GISAID identifier: EPI_ISL_2497905) and B.1 (GISAID identifier: EPL_ISL_825084) were used for the study. The isolates were propagated and passage twice in VeroCCL81 cells and sequence verified by Next generation sequencing. The virus stock was titrated to determine the 50% tissue culture infective dose (TCID50)/ml.

### Animal experiments

For the virus shedding study, eighteen female, Syrian hamsters of 8-10-week-old age were divided into 3 groups of 6 animals each. Six hamsters each were challenged with 0.1 ml of 10^5^ TCID50/ml of B.1/Delta/B.1.617.3 virus variants intranasally under isoflurane anesthesia. The animals were monitored for any clinical signs and the body weight loss. Throat swabs (TS), nasal wash (NW) and faecal samples were collected in 1 ml virus transport media on alternate days till 14 days for viral load estimation. To study the pathogenicity of each variant, 3 groups of 16 female Syrian hamsters of 8-10-week-old age used. The hamsters were challenged with 0.1 ml of 10^5^ TCID50/ml of B.1/Delta/B.1.617.3 virus variants intranasally under isoflurane anesthesia. Four hamsters from each group were euthanized on 3, 5, 7- and 14-days post infection (DPI) by isoflurane anesthesia overdose to collect blood and organ samples.

### Viral RNA detection

The tissue homogenate/ swab specimens were used for RNA extraction. MagMAX™ Viral/Pathogen Nucleic Acid Isolation Kit was used for viral RNA extraction and real-time RT-PCR was performed for E gene of SARS-CoV-2 and for N gene for the sub genomic (sg) RNA detection using the published primers [13, 14].

### Anti-SARS-CoV-2 antibody detection

The serum samples collected on day 3, 5, 7 and 14 DPI were tested for IgG antibodies by a hamster anti-SARS-CoV-2 IgG ELISA as described earlier [15]. Plaque reduction neutralization test (PRNT) was performed to understand the neutralization ability of the sera of variant infected hamsters with Delta, B.1.617.3, B.1 and Beta variant as described earlier [16].

### Histopathological examination

The lungs samples collected during necropsy were immersion fixed with 10% neutral buffered formalin. The tissues were processed with routine histopathological techniques and were stained by hematoxylin and eosin staining. The lesions were graded on a numerical scale from 0 to 4 as no abnormality (0), minimal (1), mild (2), moderate (3) and severe (4) based on the severity of lesions observed by blinded scoring. Lesions graded include vascular inflammatory changes like congestion, hemorrhages, perivascular and peribronchial mononuclear cellular infiltration, alveolar parenchymal changes, consolidation, hyaline changes, oedema, septal thickening etc.

### Statistical analysis

The data collected from the study was analysed using Graph pad Prism version 8.4.3. For statistical analysis; non parametric Mann Whitney tests were used and the p-values less than 0.05 were considered to be statistically significant.

## Results

### Clinical observations and virus shedding

The average body weight gain in hamsters was least in the Delta variant group compared to B.1 and B.1.617.3 during the first week of infection (Figure 1a). For the Delta variant, the viral gRNA could be detected in the NW and TS samples till 7 DPI whereas in few animals of B.1 and B.1.617.3 it could be detected up to 12 and 14 DPI respectively **(Figure 1b-1d)**. The viral load was higher in the TS and NW samples compared to faecal samples. The viral gRNA load was higher during the first week post infection which decreased further and was detected in few animals only post first week in B.1 and B.1.617.3 groups. Post 5 DPI, sgRNA could be detected only in 1/6 animals in B.1 and 2/6 animals of B.1.617.3 group and in none of the animals of Delta variant group **(Figure 1e-1f)**.

**Figure 1.**
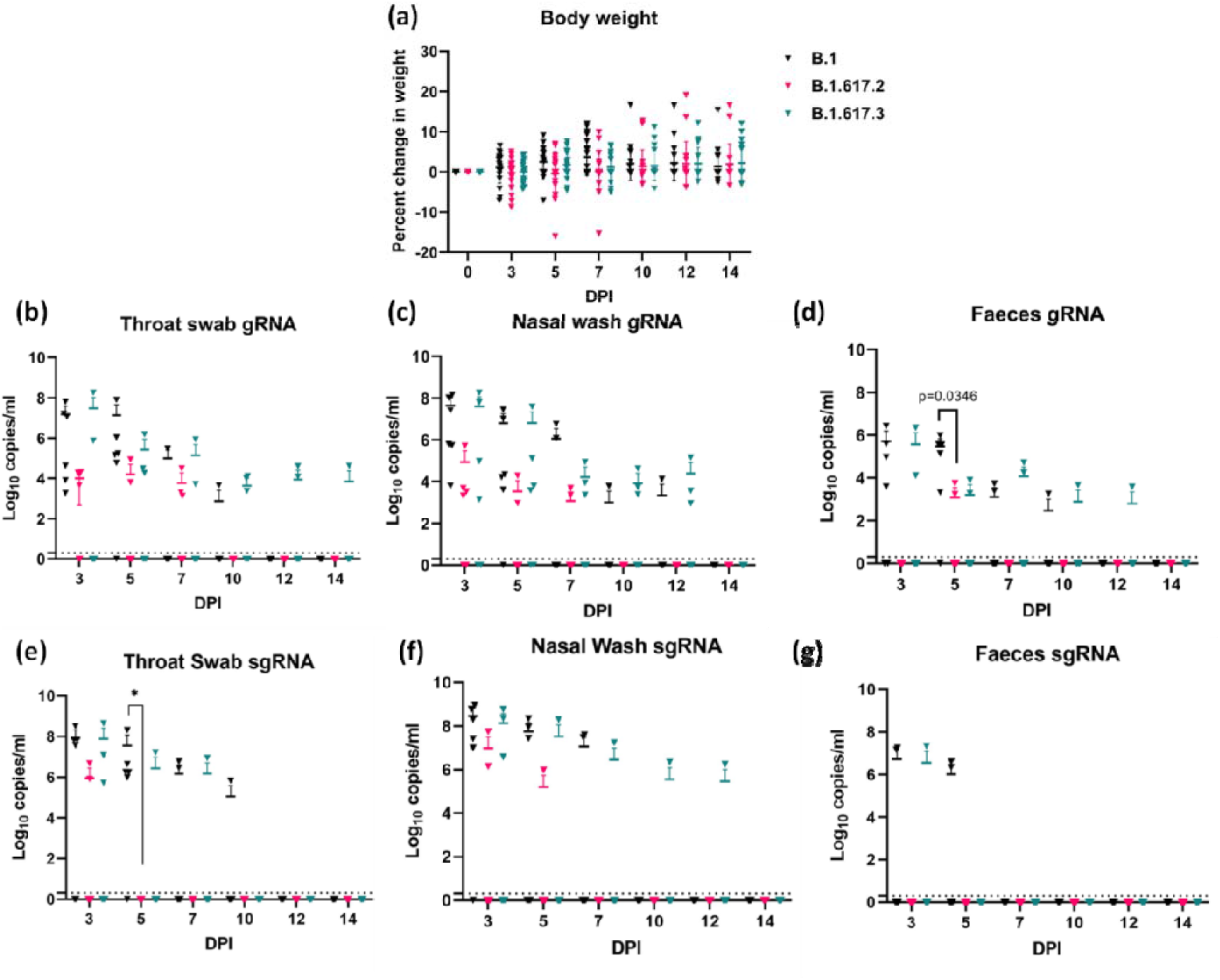
Bodyweight and virus shed by hamsters post challenge with SARS-CoV-2 variants. **(a)** Percent body weight changes in hamsters post virus infection. Viral genomic RNA load in **(b)** throat swab, **(c)** nasal wash, **(d)** faeces samples collected on 2, 4, 6, 8, 10 and 14 DPI. Viral sub genomic mRNA load in **(e)** throat swab, **(f)** nasal wash and **(g)** faeces in hamsters post virus challenge on 2, 4, 6,8,10 and 14 DPI.

### Viral load in the respiratory organs

In the nasal turbinates, trachea and lungs samples, viral gRNA could be detected till day 14 **(Figure 2a-2c)**. Nasal turbinate showed higher viral load compared to other organs in all the groups. The SARS-CoV-2 viral RNA detected in lungs, nasal turbinates and trachea of hamsters infected with different variants did not show any statistical difference when compared among the groups on day 3, 5 and 7 post infections. By day 14, viral gRNA clearance was observed in 3/4 animals in the B.1 variant infected group whereas gRNA could be detected in all animals in the Delta and B.1.617.3 group. The lung viral RNA was significantly higher (p=0.0286) in the lungs samples of B.1.617.3 variant on day 14 in comparison with B.1. Further we performed sgRNA detection in these samples, which showed detection of sgRNA in the lung and nasal turbinate samples till 7DPI in all the groups. On 14 DPI, all the hamsters of Delta variant infected group showed presence of sgRNA with a significantly high average copy of 2.1×10^7^ and 2.1×10^6^ in nasal turbinates and lungs respectively whereas in B.1 and B.1.617.3 group only ¼ hamsters showed presence of sgRNA **(Figure 2d-2f)**.

**Figure 2.**
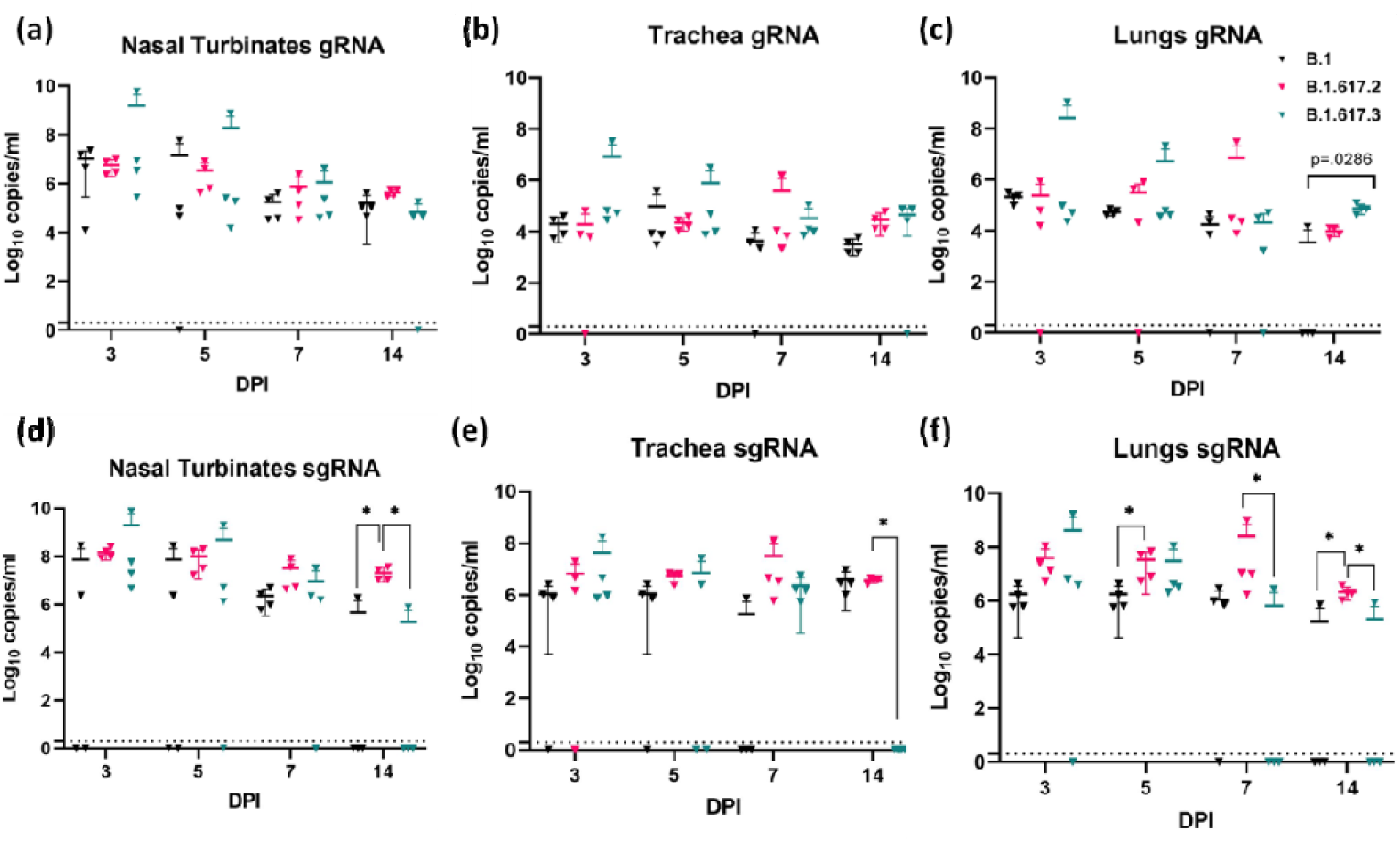
Viral load in respiratory tract samples of hamster’s post infection: Viral genomic RNA load in **(b)** nasal turbinates, **(c)** trachea, **(d)** lungs samples collected on 3, 5, 7 and 14 DPI. Viral sub genomic mRNA load in **(b)** nasal turbinates, **(c)** trachea, **(d)** lungs samples collected on 3, 5, 7 and 14 DPI. p<0.05 were considered statistically significant.

### Anti-SARS-CoV-2 immune response

Anti-SARS-CoV-2 IgG antibodies could be detected in all the animals infected from 3DPI which showed an increasing O.D value on further time points **(Figure 3a)**. The B.1 variant infected hamsters showed about 1.7-fold geometric mean titre (GMT) reduction in neutralization titre against Delta and B.1.617.3 variants and 2.3-fold reduction against the Beta variant (p<0.05). Delta and B.1.617.3 variant infected hamster sera also showed significant reduction (p<0.05) in GMT i.e., about 2.5- and 2.9-fold reduction against Beta variant. The GMT in case of Delta variant infected hamster sera was 1.8-fold reduced against the B.1 and B.1.617.3 whereas the titre of B.1.617.3 infected sera was found to be 1.1 fold reduced against Delta and 1.3 fold reduced against B.1.

**Figure 3.**
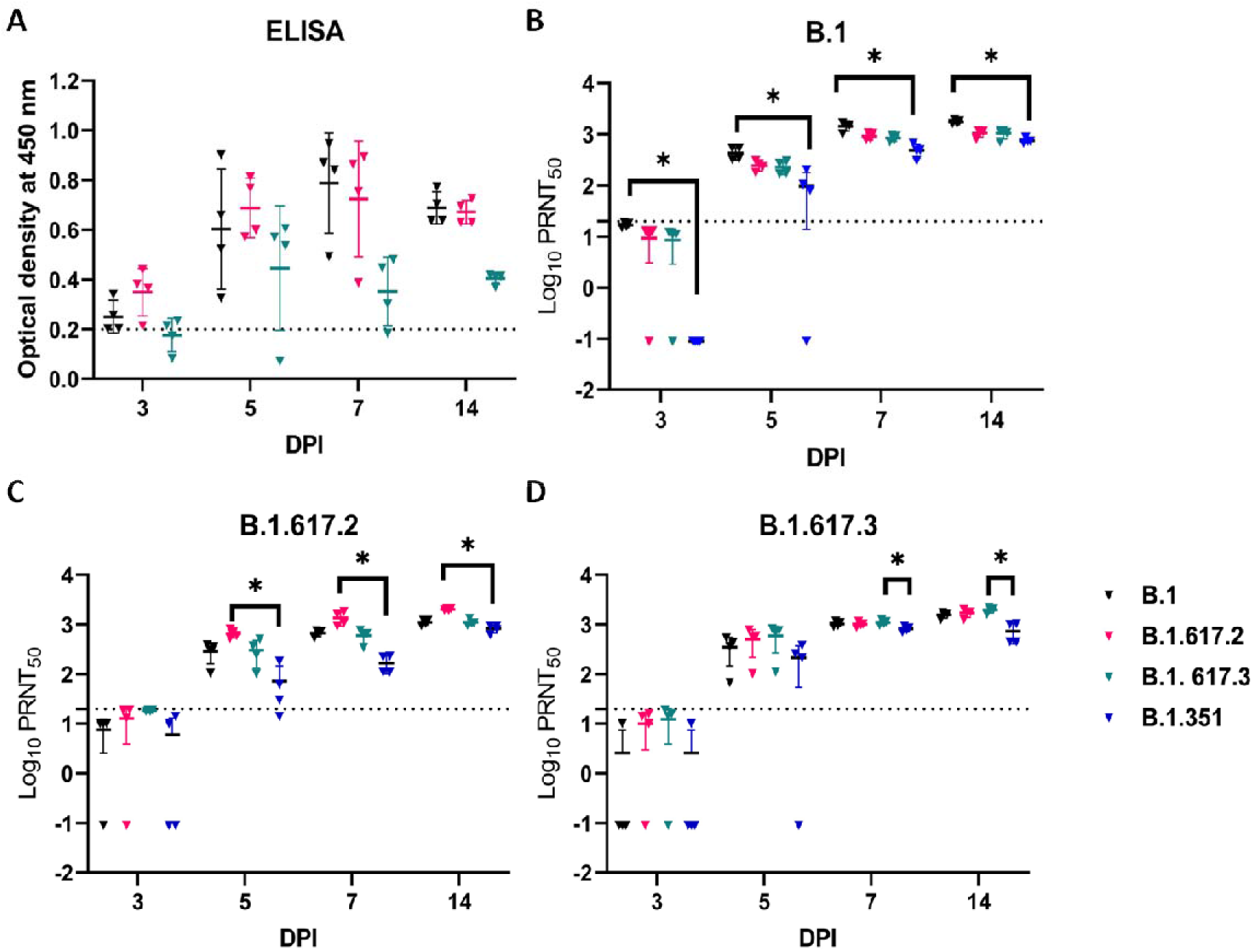
Antibody response in hamsters post challenge with SARS-CoV-2 variants: **(a)** Anti-SARS-CoV-2 IgG response in hamsters post virus infection by ELISA. Neutralizing antibody response in hamsters infected with **(b)** B.1 **(c)** B.1.617.2 (Delta) and **(d)** B.1.617.3 variants against B.1, Delta, B.1.617.3 and B.1.351 (Beta). The dotted line indicates the limit of detection of the assay.

### Lung pathology in infected hamsters

Grossly 2/16 of B.1, 6/16 of B.1.617.2 and 3/ 16 of B.1.617.3 infected hamsters showed congestion and hemorrhages. A slightly higher mean lungs weight to body weight ratio was observed on day 5 and 7 in the B.1.617.2 infected hamsters **(Figure 4a)**. The cumulative lung histopathological score also showed delta variant infected animals with a higher average score during first week of infection **(Figure 4b)**. Three/nine of the Delta and 1/9 of the B.1.617.3 infected hamsters sacrificed during the first week of infection showed diffuse areas of consolidation, hemorrhages, pneumocyte hyperplasia, septal thickening and perivascular and peribronchial inflammatory cell infiltration of moderate severity and one hamster of B.1.617.2 group showed severe lesions **(Figure 5a-d)**. Lung tissues showed minimal to mild pathological changes in case of all the B.1 variant infected hamsters. The lesions observed were mostly focal on all days of sacrifice except in 2/4 hamsters sacrificed on both day 5 and 7 which showed multifocal areas of consolidation and mononuclear cell infiltration **(Figure 5e, 5f)**.

**Figure 4.**
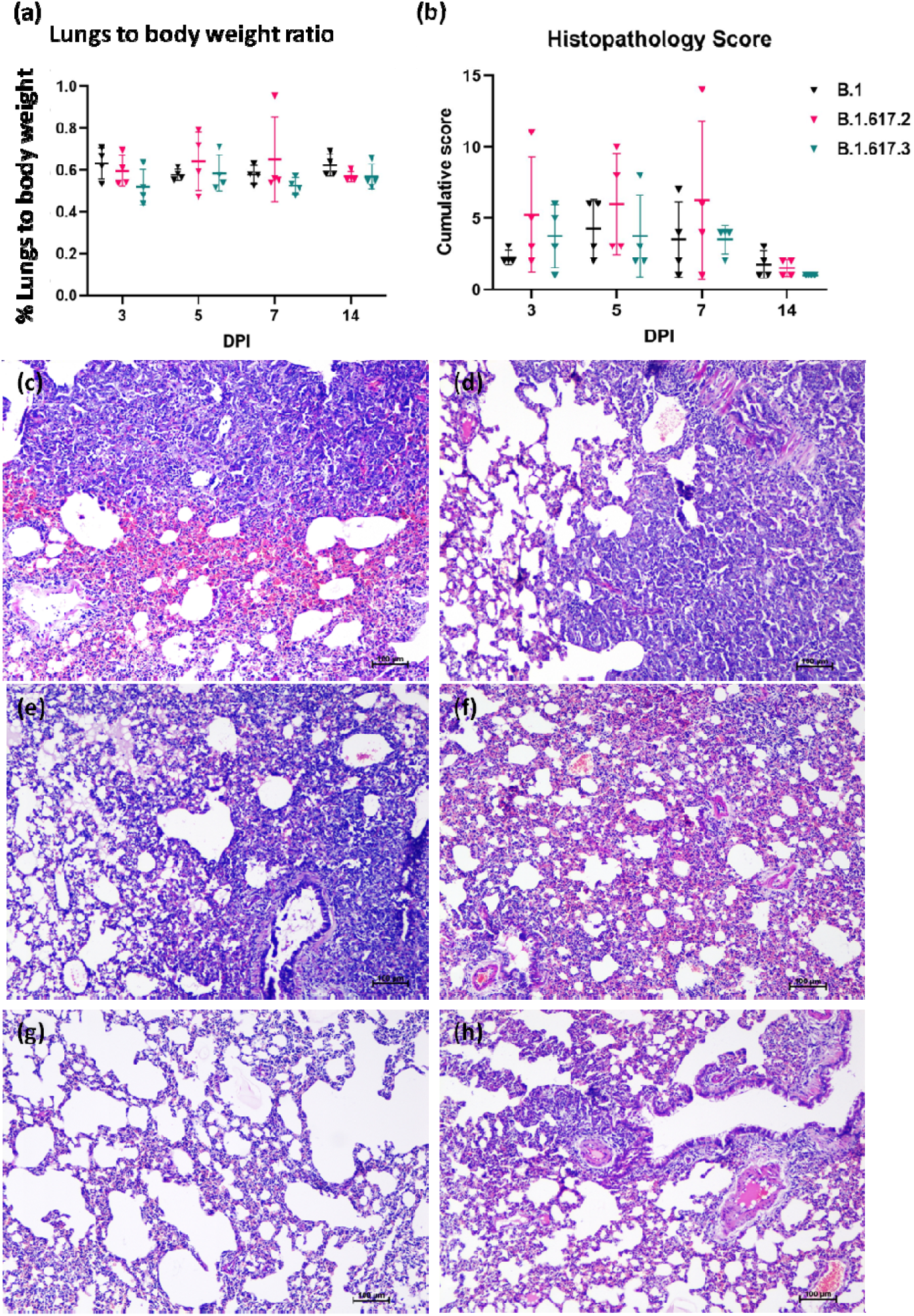
Pathological changes in lungs observed at necropsy in hamsters post infection with B.1, Delta and B.1.617.3 variants: **(a)** Proportion of lungs weight to body weight of hamsters at necropsy. **(b)** Cumulative lung histopathology score in hamsters infected with SARS-CoV-2 variants. Lungs of Delta variant infected hamsters showing **(c)** diffuse congestion and hemorrhages in the lung parenchyma **(d)** foci of diffuse mononuclear infiltration and consolidation with mild congestion. Lungs of B.1.617.3 infected hamster showing **(e)** alveolar septal thickening, exudation and hyaline changes in the alveoli **(f)** congestion and alveolar septal thickening. Lungs of B.1 infected hamster showing **(g)** few small foci of mononuclear cell infiltration and **(h)** congestion and perivascular inflammatory cell infiltration.

## Discussion

Delta variant possess SARS-CoV-2 spike protein mutations which are known to affect virus transmissibility and neutralization. The secondary attack rate of the variant is also higher compared to the Alpha variant [7]. The variant is spreading faster across the globe to become dominant variant in many countries [1]. Virus shedding from an infected host is an important determinant of virus transmission. We found a higher viral load in the throat and nasal swab samples in the first week compared to faecal samples indicating the respiratory tract tropism. The earlier studies on various SARS-CoV-2 isolates of different lineages have also reported high viral loads in the initial week of infection in humans [17]. But prolonged shedding of the virus could not be observed with any of the variants studied here as reported in human cases [18]. Surprisingly, we observed the gRNA and sg mRNA in the respiratory organs till 14 days. The sensitivity of detection of viral sgRNA by nasal wash sampling post 1 week when viral gRNA load declines is less as per our observations in earlier studies in Syrian hamsters [19,20,21,22].

The prolonged detection of sg mRNA in the nasal turbinates and lungs of B.1.617.2 could be a contributing factor to support the increased transmissibility attributed to this variant. Viral sg mRNA is considered to be an indicator of active infection [17]. In human COVID-19 cases, where presence of sgRNA is studied, it is found that its rarely detectable post 8 days of illness [23,24]. A study from England has shown increased household transmission associated with the Delta compared to the Alpha, which was reported to be highly transmissible earlier [25]. A recent modeling study from India during second COVID-19 wave has also shown the increased transmissibility and immune escape has led to the sudden rise in COVID-19 cases due to Delta [26].

The live virus neutralization assay showed a significant reduction in neutralization in case of the Beta variant with all the variants studied. Beta variant is known for reduced neutralization by many monoclonal antibodies and convalescent sera from patients infected with early SARS-CoV-2 isolates [26,27]. Although not significant, reduction was seen in case of B.1 infected hamster sera with Delta variant also. Mlcochova et al., 2021 reported 20 to 55% immune evasion by Delta variant in case of prior infections with non-B.1.617.2 lineages [26]. B.1.617.3 possesses the E484Q mutation which is a known site in RBD which can impact the serum neutralization efficiency [6]. But we found the B.1.617.3 infected animal sera better neutralizing the B.1 and Delta variants. Few recent studies reported reduced neutralization of Delta variant by the BNT162b2 mRNA vaccine [8,9]. Another study reported only modest differences in the protective antibody titres against Delta variant following two doses of the BNT162b2 or ChAdOx1 vaccine [10]. B.1.617 was found resistant to certain monoclonal antibodies approved for COVID-19 treatment like Bamlanivimab and REGN10933 [11,12]. In India, sera neutralization studies on Covishield and Covaxin vaccine recipients showed neutralization potential against B.1.617.1 and B.1.617.2 respectively [28,29].

Lung disease with severe lesions was observed in case of some Delta variant infected hamsters, indicating the potential of the variant to cause severe disease. Delta variant has shown increased replication and enhanced entry efficiency in in vitro experiments [23]. An increased fitness advantage of the variant has been observed in respiratory organoid system compared to the wild type SARS-CoV-2 [23]. Reports from England and Scotland have shown an increased risk of hospitalization in case of the B.1.617.2 variant cases [7]. Our earlier studies on pathogenicity of B.1.617.1, a variant of same lineage in hamster model have shown severe lung disease [20].

In the present study we observed no significant difference in viral RNA shedding among the different variants studied. sgRNA detection in the respiratory tract of hamsters infected with Delta variant for 14 days indicates the capability of prolonged period of persistence of variant in target organs which could be a contributing factor for its increased transmission. The neutralizing capability of the B.1, Delta and B.1.617.3 variant infected animals were found considerably lower with the Beta variant. The sera of B.1 infected animal also showed reduction in neutralizing antibodies against Delta variant. Delta variant could induce lung disease of severity in 40 % of infected animals.

## Author Contributions

Conceptualization, P.D.Y and S.M.; methodology, P.D.Y, S.M, A.S.A, G.S; software, D.N.; validation, P.D.Y, A.S.A, ; formal analysis, D.N and S.M.; investigation, S.M; resources, P.D.Y, G.S and K.L; data curation, P.D.Y; writing—original draft preparation, S.M.; writing—review and editing, P.D.Y.,N.G; visualization, P.D.Y; supervision, P.D.Y; project administration, P.D.Y.; funding acquisition, P.D.Y, N.G. All authors have read and agreed to the published version of the manuscript.

## Institutional Review Board Statement

The study was approved by the Institutional Animal Ethics Committee, ICMR-National Institute of Virology, Pune (Approval no.: NIV/IAEC/2021/MCL/01) and all the experiments were performed as per the guidelines of the Committee for the Purpose of Control and Supervision of Experiments in Animals, Government of India.

## Data Availability Statement

All the data pertaining to the study are available in the manuscript.

## Acknowledgments

Authors gratefully acknowledge the encouragement and support extended by Prof. (Dr) Balram Bhargava, Secretary to the Government of India Department of Health Research, Ministry of Health & Family Welfare & Director-General, ICMR; Dr. Sameeran Panda, Head of Epidemiology and Communicable Diseases, ICMR and Prof. Priya Abraham, Director, ICMR-National Institute of Virology, Pune. Authors are thankful to Mr. Manoj Kadam, Mr. Abhimanyu Kumar, Mr. Annasaheb Suryawanshi, Dr. Gururaj Deshpande, Dr Chandrasekhar Mote, Mr. Rajen Lakra, Dr. Rajlaxmi Jain, Mr. Prasad Sarkale, Mr. Shreekant Baradkar, Mr. Chetan Patil, Mrs. Asha Salunke, Mrs. Supriya Hundekar, Ms. Pranita Gawande, Mrs. Ashwini Waghmare, Mrs. Kaumudi Kalele, Mr. Kundan Wakchaure, Ms. Shilpa Ray, Ms. Poonam Bodake and Mrs. Priyanka Waghmare of ICMR-NIV, Pune.

